# Nuclear Pore Complexes in Various States of HL-60/S4 Cells

**DOI:** 10.64898/2026.01.09.697371

**Authors:** Ada L. Olins, Igor Prudovsky, Donald E. Olins

## Abstract

This study is focused upon how the structure and function of interphase nuclear pore complexes (NPCs) respond to various cellular stresses (i.e., cell differentiation, knockdown of Lamin B Receptor [LBR], and cellular dehydration) in a myeloid cell line (HL-60/S4). Each cellular stress was examined by GSEA (Gene Set Enrichment Analysis) to determine how the structure and function of the NPCs were affected. Cell differentiation into granulocytes and into macrophages resulted in widespread decreases in nucleoporin (NUP) transcription levels, affecting NPC structure and NPC transport capability. LBR knockdown (HL-60/sh1 cells) in undifferentiated cells exhibited major increases in NUP transcription, combined with improved NPC structural quality and transport capability, implying that nuclear pore function is not adversely affected by the loss of LBR. In contrast, cell dehydration of the undifferentiated HL-60/S4 cells in hyperosmotic culture medium resulted in disorganized nucleoporin transcription, with evidence of abnormal NPC structure and transport capability. The structural integrity and transport function of nuclear pores are clearly responsive to the various cellular stresses. Future investigations should examine the reversibility and resilience of these cells to such stresses as those described in this study.

## Introduction

Nuclear pore complexes (NPCs) embedded within the interphase nuclear envelope both influence and respond to on-going cellular physiology. NPCs are the conduits for mRNA and ribosomal RNA to exit the interphase nucleus and enter the cytoplasm. In addition, NPCs are the conduits for transcription factors, newly synthesized chromatin structural proteins and critical enzymes to enter the nucleus. Mammalian NPC molecular structure has been well described, especially employing cryo-electron microscopy (Guglielmi et al., 2020; Lin and Hoelz, 2019; Petrovic et al., 2025; Taniguchi et al., 2025). The human NPC is disc-shaped with an outer diameter of ∼120 nm, an inner channel diameter of ∼42.5 nm, a height (thickness) of ∼80 nm and a molecular mass of ∼110 MDa. Each NPC disc normally presents an 8-fold rotational symmetry, constructed with ∼1000 protein subunits, consisting of nucleoporins (NUPs). Human NPCs have ∼30 types of NUPs. Excellent schematic diagrams of the positions of the various types of NUPs within the 3-D structure of human NPCs have been published (Buchwalter et al., 2019; Guglielmi et al., 2020; Lusk et al., 2025; Petrovic et al., 2025).

HL-60/S4 cells are derived from a human Acute Myeloid Leukemia (AML) cell line (Leung et al., 1992). HL-60/S4 cells can be differentiated into granulocytes with retinoic acid (RA) (Olins et al., 1998) and into macrophage with phorbol ester (TPA) (Olins et al., 2001), each following four days of drug exposure. Undifferentiated HL-60/S4 (0) cells exhibit robust growth in suspension with a doubling time of ∼ 17 hours. RA-treated HL-60/S4 cells exhibit a gradual slowing of cell division and remain in suspension during differentiation. RA-induced granulocytes exhibit significant nuclear shape changes during differentiation. Nuclear lobulation with multiple surface nuclear envelope chromatin sheets (ELCS), characterizes the granulocyte differentiated state (Olins et al., 1998). TPA-treated cells attach to a substrate and cease division within one day. Comparative transcriptomes of undifferentiated HL-60/S4 cells and the RA and TPA-differentiated cells, after four days of drug exposure, have been published (Mark Welch et al., 2017) and are employed in this study. Comparisons of two other HL-60/S4 cell states with undifferentiated (0) cells are also included in this study: 1) Lamin B Receptor (LBR) knockdown in HL-60/sh1 0 cells, described earlier (Mark Welch et al., 2024; Olins et al., 2010; Olins et al., 2025b); 2) Hyperosmotic stressed HL-60/S4 0 cells, also described earlier (Mark Welch et al., 2022; Olins et al., 2020; Olins et al., 2025a). The Results section of this article is divided into three parts reflecting the three separate NPC “stress” comparisons: 1) RA and TPA-differentiation versus undifferentiated S4 0 cells; 2) LBR knockdown sh1 0 versus S4 0 cells; 3) Hyperosmotic, 300 mM sucrose-treated S4 0 cells, stressed for 30 and 60 minutes versus unstressed S4 0 cells.

## Materials and Methods

### Cell Cultivation

HL-60/S4 cells can be purchased from ATCC (CRL-3306). They are cultivated in RPMI 1640 medium + 10% (unheated) Fetal calf serum + 1% Pen/Strep/Glut. We employ T-25 (6 ml) or T-75 (18 ml) flasks, lying horizontal to maximize surface area. The cells were grown in a humidified incubator at 37o C with 5% CO_2_. HL-60/sh1 and HL-60/gfp were generated and cultivated as described (Olins et al., 2010a). Unfortunately, the HL-60/gfp culture was lost in a laboratory accident. However, this cell loss occurred after undifferentiated and differentiated mRNA transcriptomes were generated. The HL-60/sh1 and HL-60/gfp cell lines were cultivated in normal growth medium plus 1μl/ml puromycin (1:1000 dilution of puromycin stock), to maintain selection pressure on the lentivirus infected cell lines. Puromycin was not present during cell differentiation.

### Nuclear Pore Gene Clusters and Gene Names

There are several different published sets of nomenclature for the nuclear pore structure and gene names. In this paper, we base our nomenclature upon an earlier version (Buchwalter et al., 2019). Major differences are highlighted in Table 1. The earlier version of NPC nomenclature is utilized in Figures 2, 8 and 13 of this study.

**Table 1.** Differences in Nomenclature, comparing the present study with a published review (Petrovic et al., 2025) on the location of NUPs within nuclear pore complexes (NPC).

| **NOMENCLATURE** |  |
| --- | --- |
| **Figure 2 (this paper)** | **Figure 1 (Petrovic et al., 2025)** |
| **GENE CLUSTERS** |  |
| Cytoplasmic Filaments | Cytoplasmic filaments |
| Nuclear Ring |  |
| Central Channel | Inner ring nups |
| Central Pore |  |
| Cytoplasmic Ring | Coat nups |
| Nuclear Envelope | POMs |
| Nuclear Basket | Nuclear basket |

| GENE NAMES |  |
| --- | --- |
| RANBP2 | NUP358 |
| NUP35 | NUP53 |
| NUP85 | NUP75 |
| NUP98 | NUP96 |
| AHCTF1 | ELYS |
| NDC1 | ALADIN |

**Figure 1.**
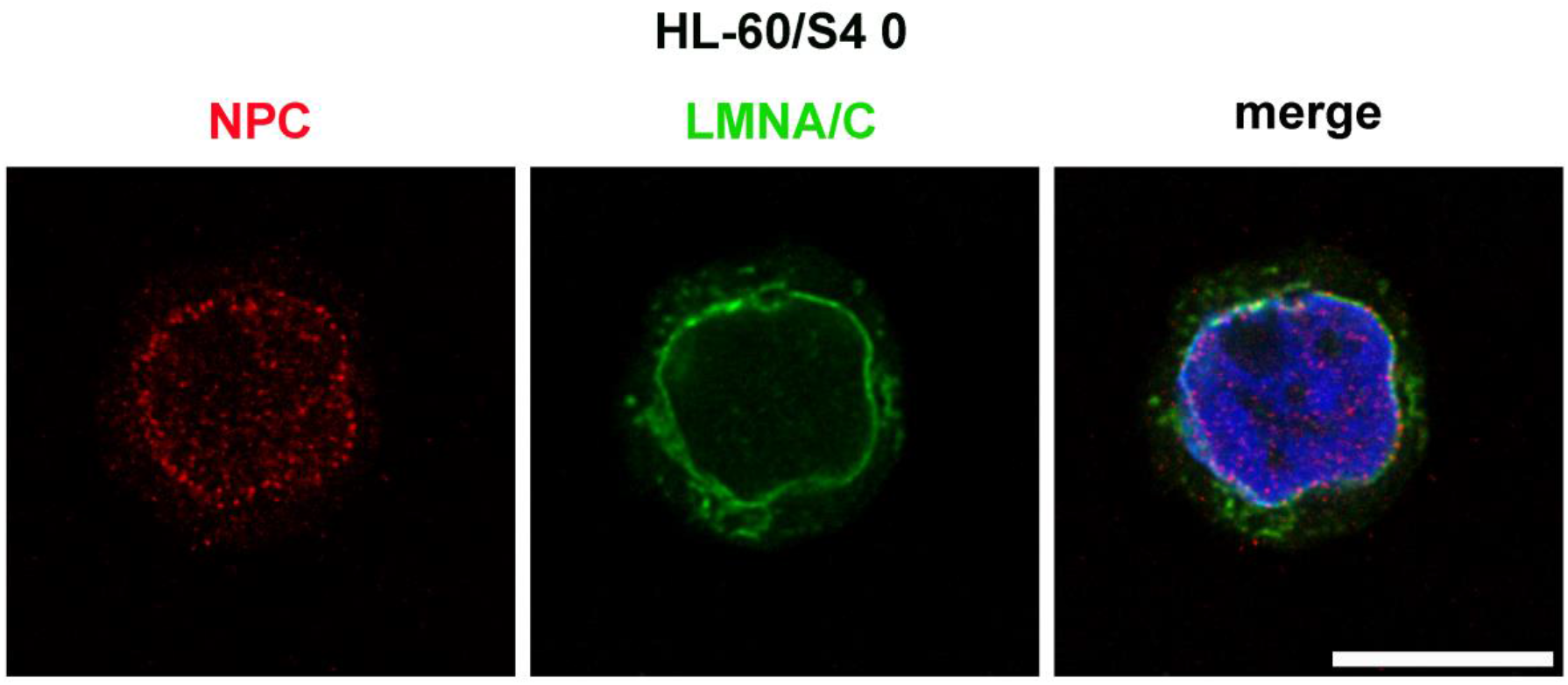
Immunostaining of an undifferentiated HL-60/S4 cells with two different antibodies: Mouse anti-nuclear pores complexes (Mab 414), Red; Rabbit anti-Lamin A/C, Green, combined with DAPI, Blue. Note the enrichment of NPCs in the vicinity of the nuclear envelope, which is highlighted by Lamin A/C staining. All images are deconvolved. The magnification bar represents 10 μm.

**Figure 2.**
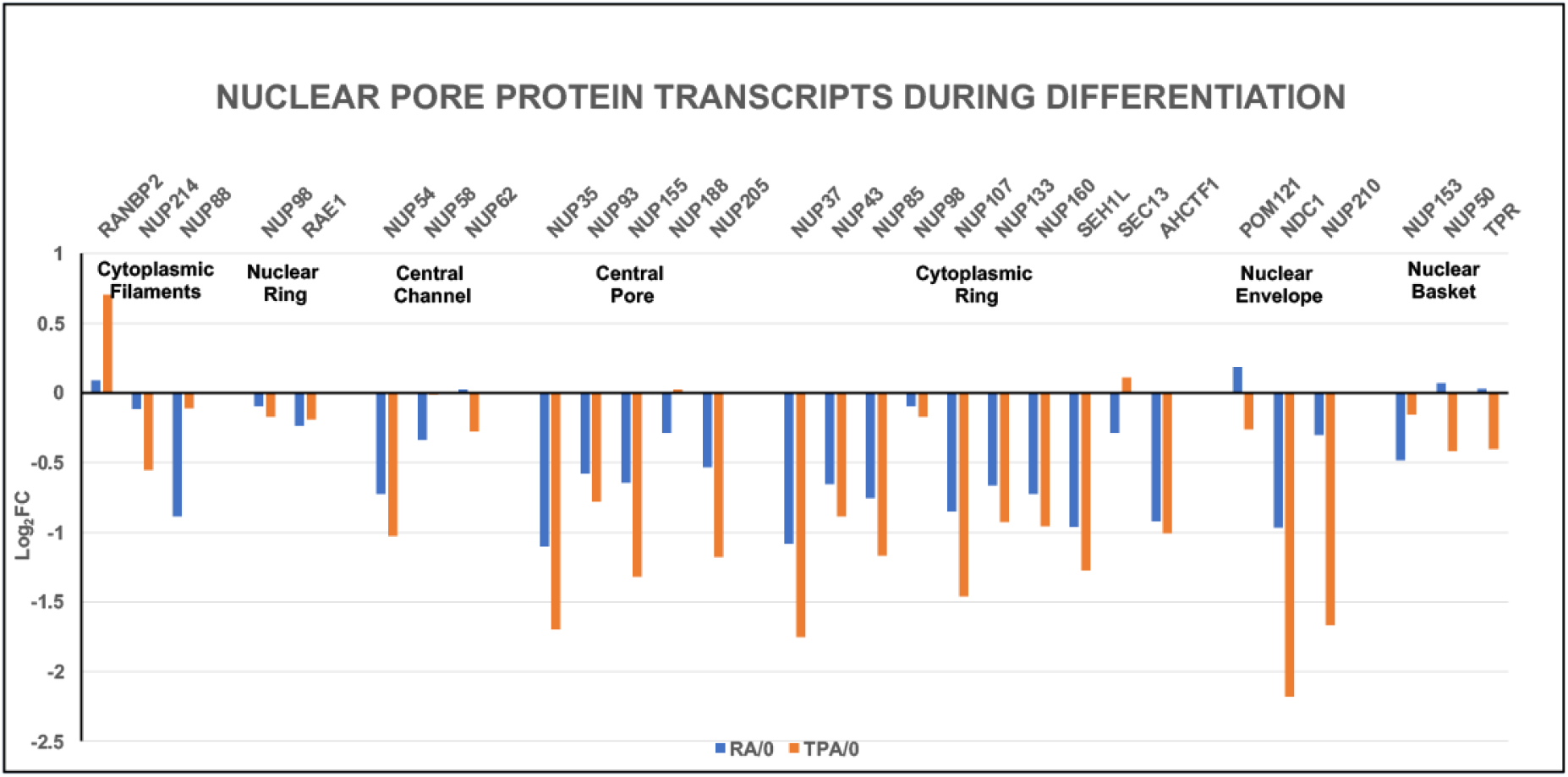
Nuclear pore protein relative transcript levels (Log_2_FC) following cell differentiation. Cell states: RA/0, granulocytes compared to undifferentiated HL-60/S4 cells; TPA/0, macrophages compared to undifferentiated HL-60/S4 cells. Seven NPC structural regions are indicated, with resident gene names clustered together. A recent reclassification of NPC structural regions with resident clustered gene names has been published (Petrovic et al., 2025). See Table 1 for a summary of the differences between some gene clusters and some gene names compared to those employed in the present study.

### Immunostaining

NPCs were visualized with Mouse monoclonal Mab 414 (Abcam 24609), which recognizes several NUPs (RANBP2, NUP214, NUP153 and NUP62). Cells were attached to polylysine-coated slides by settling for 30 minutes, fixed for 15 minutes with 3.7% HCHO/0.5xPBS, washed with PBS, permeabilized with 0.1% Triton-X100/PBS for 30 minutes, blocked with 5% normal goat serum/PBS for 30 min and stained with Mab 414 (1:200) for 1 hour, following procedures described earlier (Mark Welch et al., 2024; Olins et al., 2025a; Olins et al., 2025b). Table 2 lists the two antibodies used in Figure 1, their sources and dilutions employed for immunostaining. Figure 1 displays immunostaining of an undifferentiated HL-60/S4 cell, demonstrating that most of the NPCs are confined to the interphase nuclear envelope, with some of NUP epitopes dispersed within the nucleus. These scattered dots may represent NUPs that are not integrated into mature NPCs.

**Table 2.**
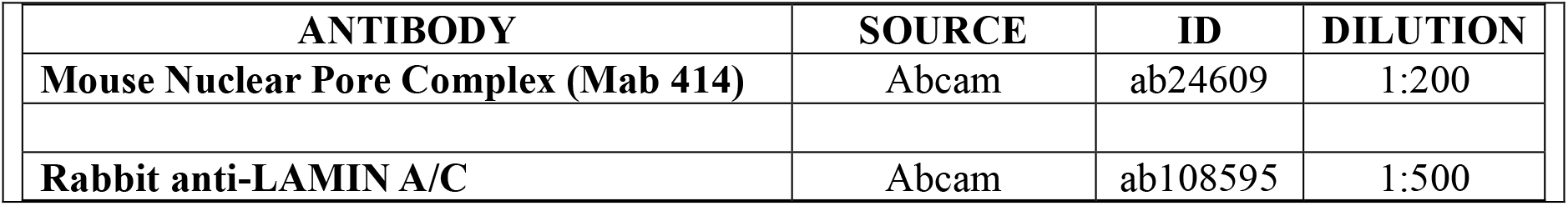
Antibodies, Sources and Dilutions.

| ANTIBODY | SOURCE | ID | DILUTION |
| --- | --- | --- | --- |
| Mouse Nuclear Pore Complex (Mab 414) | Abcam | ab24609 | 1:200 |
| Rabbit anti-LAMIN A/C | Abcam | ab108595 | 1:500 |

### GSEA Analyses: Three sets of transcriptomic data were analyzed in this study: Cell Differentiation; Loss of LBR and Cell Dehydration

The data, formatted for direct insertion into GSEA as a data set, which can be probed with your favorite Gene-set, are located in the Supplemental Data Tables S1, S2 and S3. Table S1 contains the expression data for HL-60/S4 cells differentiated with Retinoic Acid (RA), Phorbol Ester (TPA) and for untreated cells. Table S2 is the data for cells in which LBR has been knocked down. Table S3 is the expression data for cells treated with 300 mM sucrose for 0, 30, and 60 minutes.

## Results

### Part I. The stress of cell differentiation

#### Differentiation of HL-60/S4 cells produces downregulation of NUP protein transcripts and likely malfunction of NPC transport

A graph of the relative mRNA levels (Log_2_FC) of 29 NUP structural protein transcripts at 4 days of RA and TPA differentiation of HL-60/S4 cells is shown in Figure 2. The transcripts are assigned to 7 structural regions (Gene Clusters) of the NPC, based upon published diagrams (Buchwalter et al., 2019; Guglielmi et al., 2020). With a few exceptions, the impression from Figure 2 is that there is a significant downregulation of most NUP structural protein transcripts during **both** RA and TPA differentiation in all 7 NPC structural regions.

#### Importins, Transportins and Exportins

These transport proteins carry various cargoes into and out of cell nuclei, from and to the cytoplasm (Yang et al., 2023). Figure 3 convincingly demonstrates that the transport protein transcripts are also significantly downregulated during differentiation induced by RA and by TPA. Combined with the data on NUP transcripts, a significant decrease of normal nuclear pore activity may occur during cell differentiation.

**Figure 3.**
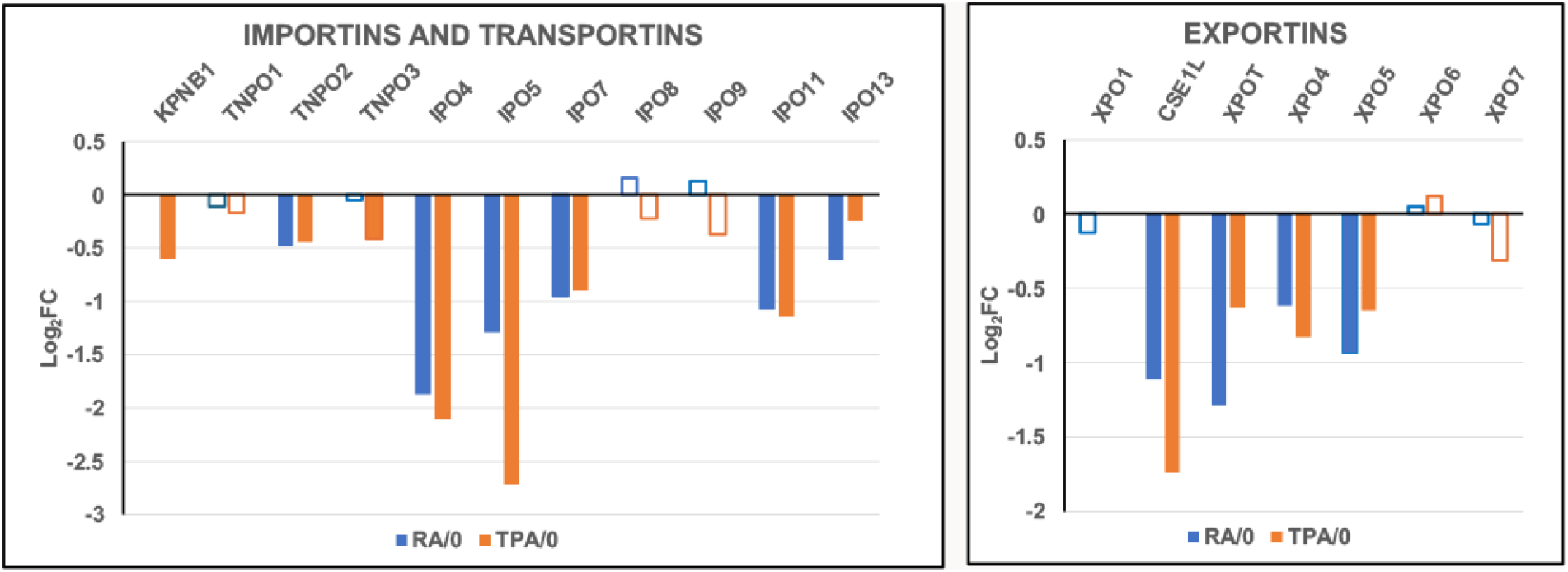
Importins, Transportins and Exportins relative transcript levels (Log_2_FC) following cell differentiation. Cell states: RA/0, granulocytes compared to undifferentiated HL-60/S4 cells; TPA/0, macrophages compared to undifferentiated HL-60/S4 cells. The open bars indicate lower statistical significance (PPDE <0.95); solid bars indicate higher statistical significance (PPDE > 0.95).

Gene Set Enrichment Analysis (GSEA) was performed to examine the predicted structural and functional quality of the differentiated nuclear pores following RA and TPA treatment for 4 days (Figure 4). [For a detailed description of GSEA plot parameters, see (Olins et al., 2025b)]. Four gene sets are illustrated in both RA and TPA treatment, with the plot parameters shown in Table 3. The negative NES (Normalized Enrichment Score) exhibit generally significant P values (Nominal p). These GSEA plots argue that the NPC structures may be malformed, possibly missing the nuclear basket and the outer ring. Figure 5 and Table 4 also support that Nuclear Transport is adversely affected by differentiation of the HL-60/S4 cells. Taken together, the graphical presentations of the NPC relative transcription changes and the GSEA NPC structural and functional analyses imply that the RA and TPA-treated cells are declining in their transport capabilities compared to the rapidly growing undifferentiated HL-60/S4 cells.

**Table 3.**
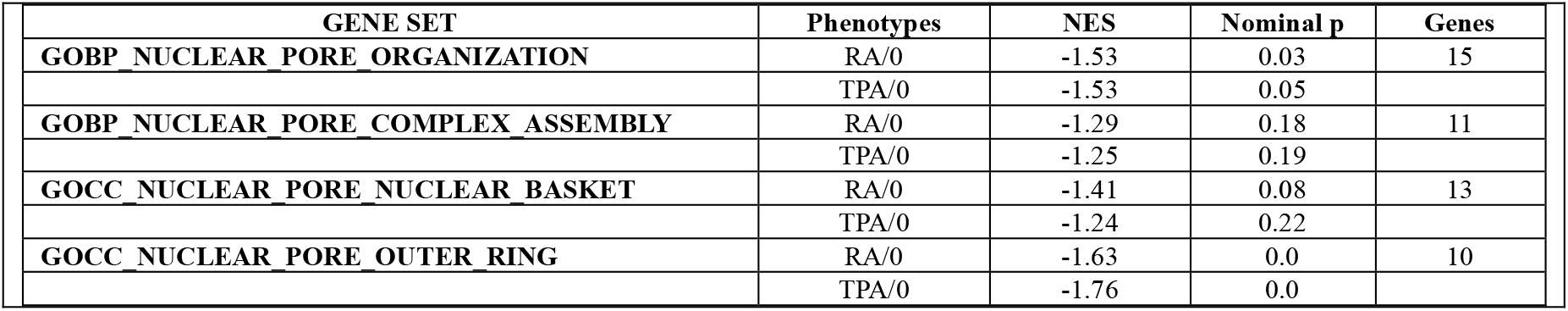
GSEA plot parameters from Figure 4. The negative NES values indicate the relative paucity of gene set transcripts in the differentiated phenotype. The Nominal p values indicate that most of the NES scores (and their implications) are quite significant.

| GENE SET | Phenotypes | NES | Nominal p | Genes |
| --- | --- | --- | --- | --- |
| GOBP_NUCLEAR_PORE_ORGANIZATION | RA/0 | -1.53 | 0.03 | 15 |
|  | TPA/0 | -1.53 | 0.05 |  |
| GOBP_NUCLEAR_PORE_COMPLEX_ASSEMBLY | RA/0 | -1.29 | 0.18 | 11 |
|  | TPA/0 | -1.25 | 0.19 |  |
| GOCC_NUCLEAR_PORE_NUCLEAR_BASKET | RA/0 | -1.41 | 0.08 | 13 |
|  | TPA/0 | -1.24 | 0.22 |  |
| GOCC_NUCLEAR_PORE_OUTER_RING | RA/0 | -1.63 | 0.0 | 10 |
|  | TPA/0 | -1.76 | 0.0 |  |

**Table 4.** GSEA plot parameters from Figure 5. The negative NES values indicate the relative paucity of gene set transcripts in the differentiated phenotype. The Nominal p values indicate that most of the NES scores and their functional implications are quite significant.

| GENE SET | Phenotypes | NES | Nominal p | Genes |
| --- | --- | --- | --- | --- |
| GOMF_NUCLEAR_EXPORT_SIGNAL_RECEPTOR_ACTIVITY | RA/0 | -1.38 | 0.10 | 12 |
|  | TPA/0 | -1.51 | 0.05 |  |
| GOMF_NUCLEAR_IMPORT_SIGNAL_RECEPTOR_ACTIVITY | RA/0 | -1.53 | 0.01 | 20 |
|  | TPA/0 | -1.49 | 0.03 |  |
| GOBP_NUCLEAR_EXPORT | RA/0 | -1.21 | 0.04 | 169 |
|  | TPA/0 | -1.19 | 0.03 |  |
| GOBP_NUCLEAR_TRANSPORT | RA/0 | -1.19 | 0.04 | 331 |
|  | TPA/0 | -1.33 | 0.006 |  |

**Figure 4.**
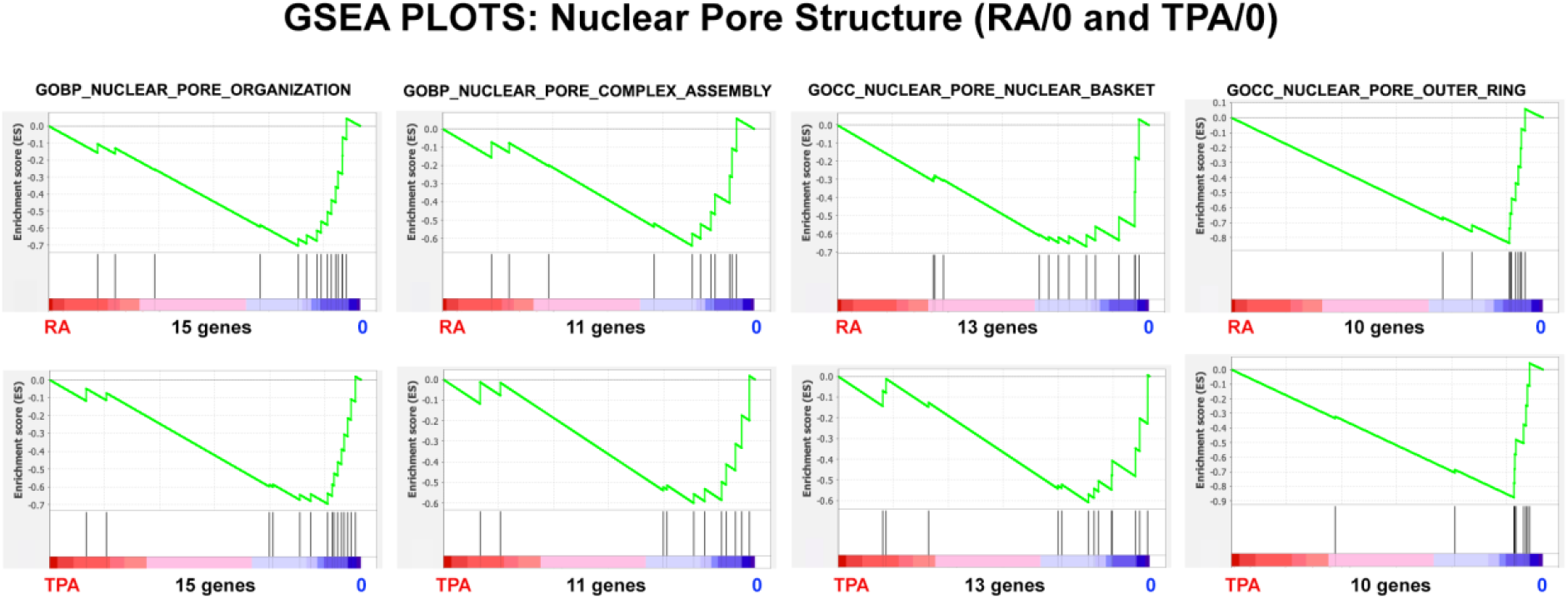
GSEA plots with four different nuclear pore structure gene sets. The number of genes in each gene set are shown at the bottom of each plot. The top row of plots compares the (RA) granulocyte phenotype to the undifferentiated cell phenotype; the bottom row compares the (TPA) macrophage phenotype to the undifferentiated cell phenotype. From the shape of these plots and the increased density of ranked genes at the undifferentiated (0) phenotype location, it is clear that most of the genes in each gene set are underrepresented in the differentiated cell phenotype region, implying that nuclear pore organization is not “normal”, possibly exhibiting structural irregularities in the two differentiated states.

**Figure 5.**
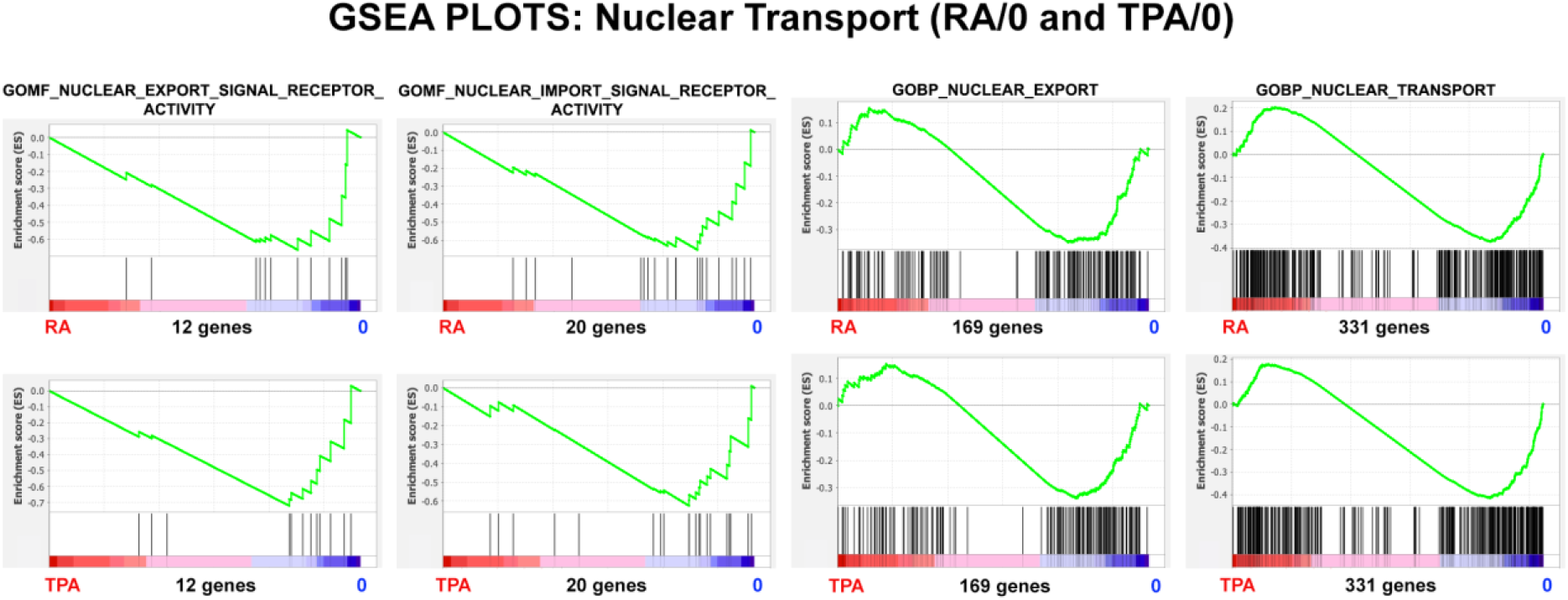
GSEA plots with four different nuclear transport gene sets. The number of genes in each gene set are shown at the bottom of each plot. The top row of plots compares the (RA) granulocyte phenotype to the undifferentiated cell phenotype; the bottom row compares the (TPA) macrophage phenotype to the undifferentiated cell phenotype. From the shape of these plots and the increased density of ranked genes at the undifferentiated (0) phenotype location, it is clear that most of the genes in each gene set are underrepresented in the differentiated cell phenotype region, implying that the nuclear pore transport function is not functioning “normally” in the differentiated states.

### HL-60/S4 RA-induced granulocytes exhibit nuclear pores within lobular nuclear envelope regions, but very seldom in ELCS

RA-induced HL-60/S4 granulocytes develop nuclear envelope (NE) heterogeneity. The NE expands its surface area, which becomes associated with additional peripheral heterochromatin during the formation of nuclear lobes and ELCS (Olins et al., 1998; Olins and Olins, 2009).

Figure 6 shows published electron micrographs illustrating nuclear pores at the surface of the nuclear lobes, but largely absent from adjacent ELCS. ELCS appear to be interrupted by “microlobulation” (i.e., small dilations in the well-defined ELCS of apposed inner nuclear membranes). The underlying mechanism for the relative absence of NUPs in ELCS is not known. However, we attribute it to the fact that the thickness of the NPC disc (∼80 nm, by cryo-electron microscopy) is greater than the space between the apposed ELCS nuclear membranes (∼60 nm, by cryo-electron microscopy) (Xu et al., 2021). In addition, ELCS are filled with two criss-cross layers of parallel ∼30 nm heterochromatin fibers (Xu et al., 2021). Recent publications have argued that heterochromatin adjacent to the inner nuclear membrane (as in ELCS) interferes with nuclear pore formation (Krull et al., 2010; McCloskey et al., 2018). NUP proteins associated with the Nuclear Basket create heterochromatin exclusion zones (HEZ), facilitating nuclear pore formation. It is of interest that formation of Nuclear Baskets appears to be reduced in RA-differentiated granulocytes (Figure 4 and Table 3), possibly increasing the incompatibility of ELCS heterochromatin with the formation of nuclear pores.

**Figure 6.**
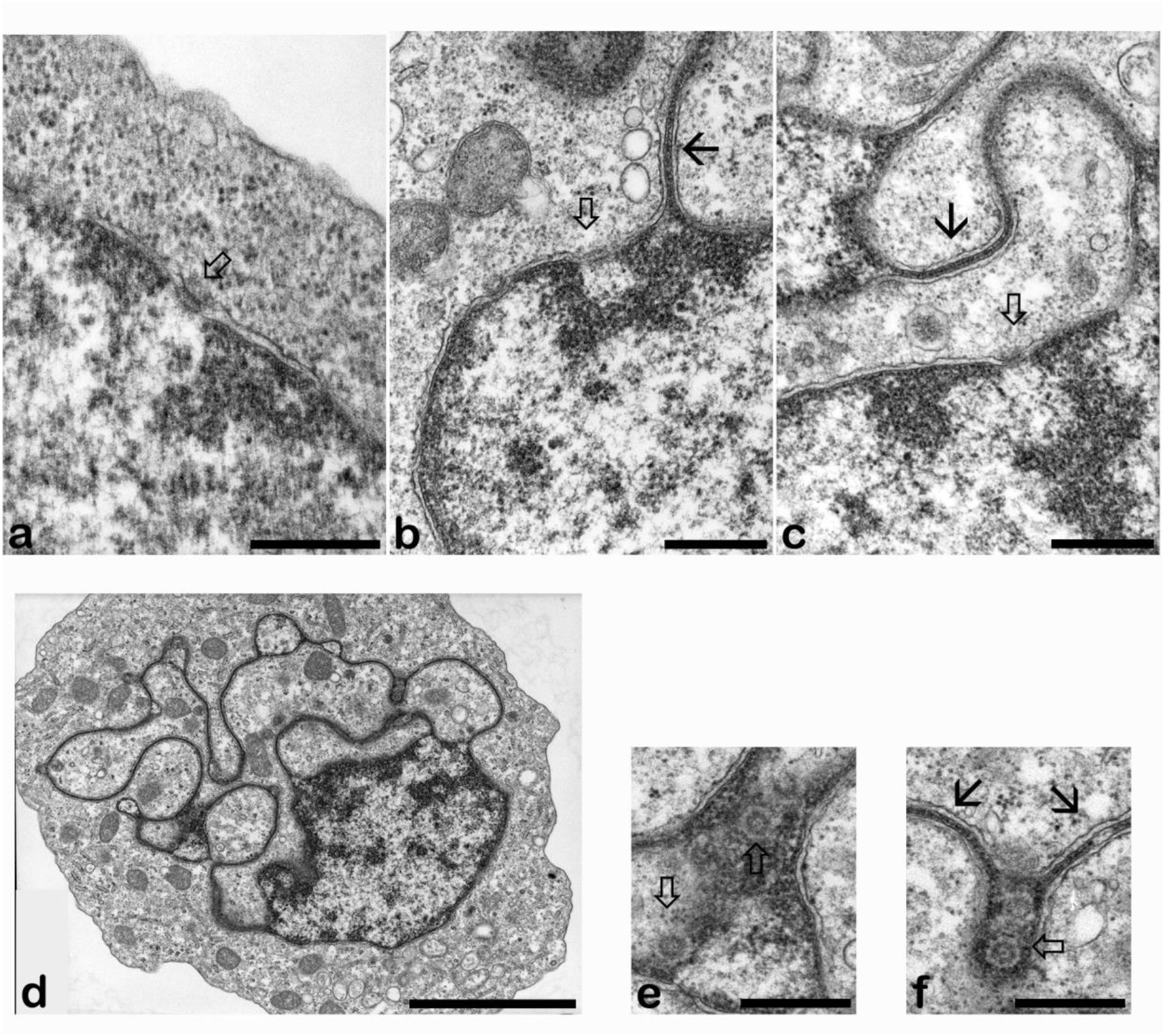
Electron microscope images of control (a) and RA-differentiated HL-60/S4 granulocytes (b, c, d, e, and f) displaying nuclear lobulation, nuclear pores and a profusion of ELCS connected to the nuclear lobes. Top Row: NPCs (open arrow heads) at the nuclear envelope of a control (undifferentiated) HL-60/S4 cell (a); Granulocytes with NPCs in nuclear lobe envelopes (b and c). ELCS are identified by thin arrows. Bottom Row: A thin section of a single HL-60/S4 granulocyte displaying a nuclear lobe affiliated with numerous ELCS (d); Two magnified regions from the same granulocyte (d) exhibit nuclear pores adjacent to surrounding ELCS (e and f) probably at the surface of microlobes. Magnification bars: a, 100 nm; b, 500 nm; c, 500 nm; d, 3 μm; e, 500 nm; f, 500 nm.

An example of Mab 414 immunostaining of an RA-differentiated HL-60/S4 granulocyte which shows four prominent nuclear lobes immunostained on their surface is presented in Figure 7. We suggest that the apparent ELCS staining arises from the interspersed “microlobes” between ELCS. Close examination of the electron micrograph (Figure 6, bottom row) indicates candidate regions of interspersed “microlobes”. Indeed, in our most comprehensive ultrastructural search for NPCs in RA-differentiated HL-60/S4 granulocytes, we stated that we “never observed nuclear pores on ELCS.” (Olins et al., 1998).

**Figure 7.**
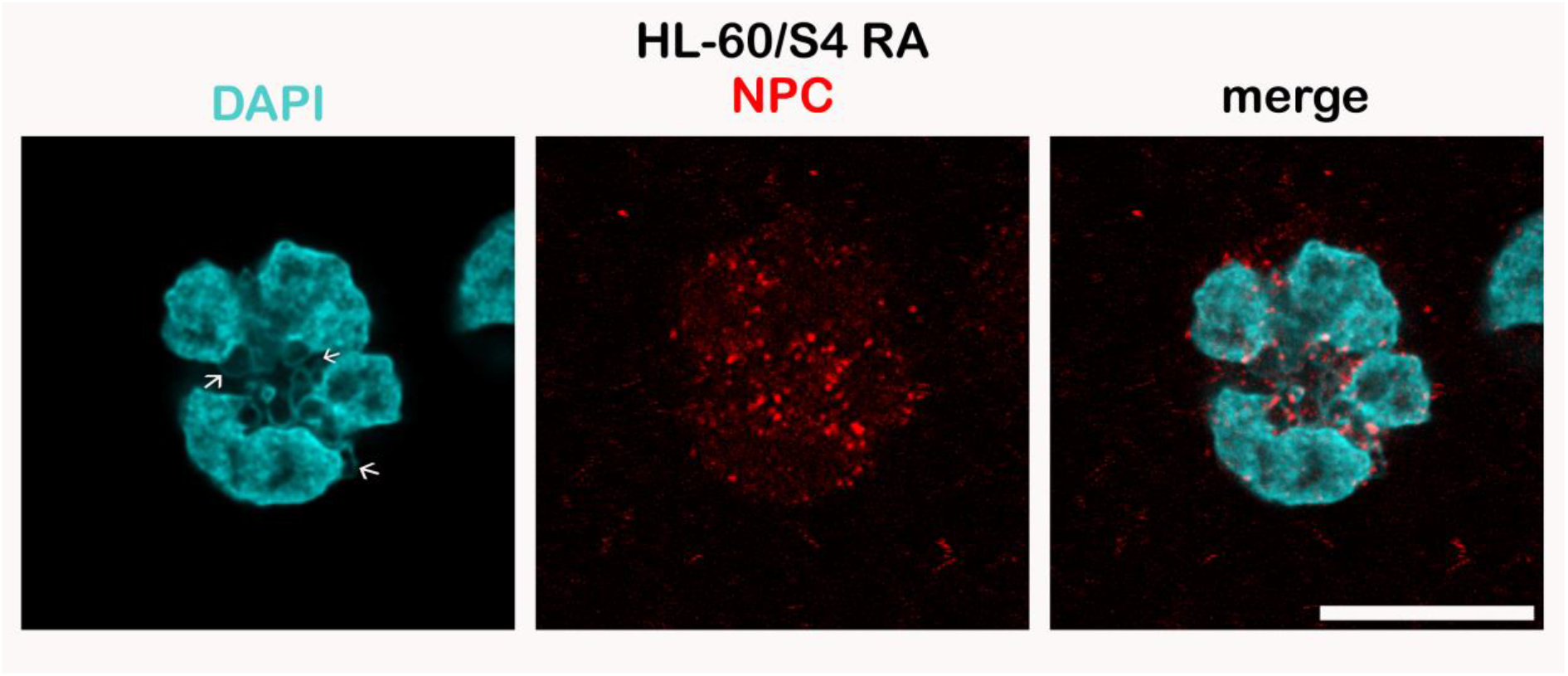
Immunostaining of a differentiated HL-60/S4 cell granulocyte with Mouse anti-nuclear pore complexes [Mab 414] (Red dots) and with DAPI (Cyan). Note the enrichment of NPCs at the surface of nuclear lobes and within the ELCS region (indicated by arrows in the DAPI image). All images are deconvolved. The magnification bar represents 10 μm.

**Figure 8.**
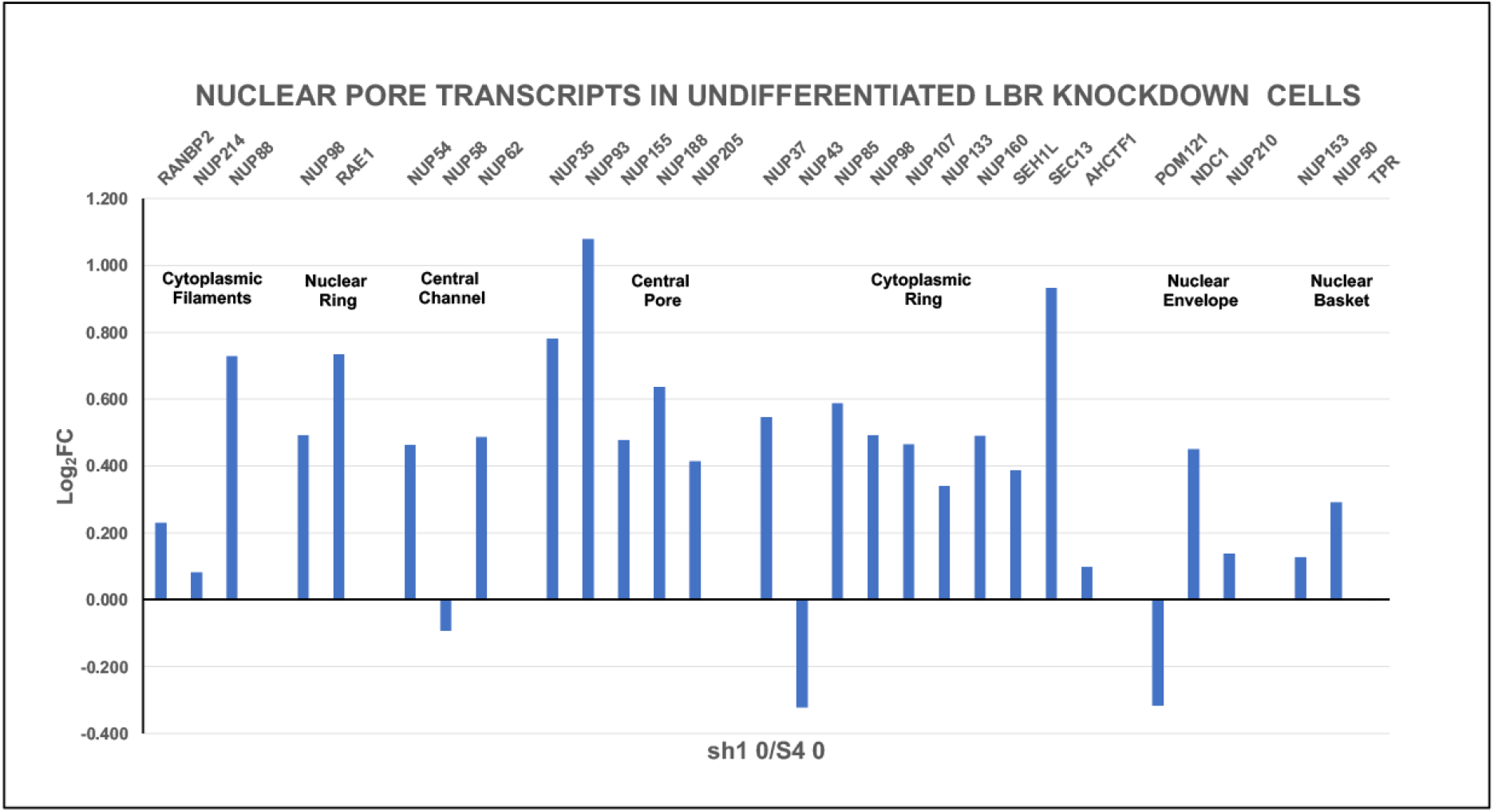
Nuclear pore protein relative transcript levels (Log_2_FC) in HL-60/sh1 0 cells, where the nuclear envelope protein LBR transcript has been knocked-down (sh1 cells). Cell states: undifferentiated (sh1 0) cells compared to undifferentiated (S4 0) HL-60/S4 cells. Seven NPC structural regions are indicated, with resident gene names clustered together as in Figure 2.

### Part II. The loss of Lamin B Receptor

#### LBR knockdown in undifferentiated HL-60/sh1 0 cells appears to upregulate NUP protein transcription

Figure 8 demonstrates that most of the 29 NUP protein transcripts are increased in the sh1 0 cells, compared to the NUP protein transcripts from control undifferentiated HL-60/S4 0 cells.

Nuclear pore immunostaining images, Mab 414, comparing undifferentiated S4 and sh1 cells are shown in Figure 9. Based upon immunostaining, there is no obvious difference between the two cell states; most of the NPCs appear located within the nuclear envelope region. We previously demonstrated that the LBR transcripts are reduced ∼8-fold in sh1 0 cells, compared to S4 0 cells (Mark Welch et al., 2024). Clearly, the NUP protein transcript changes in sh1 0 cells, Figure 8, do not resemble those seen following RA and TPA induced cell differentiation Figure 2.

**Figure 9.**
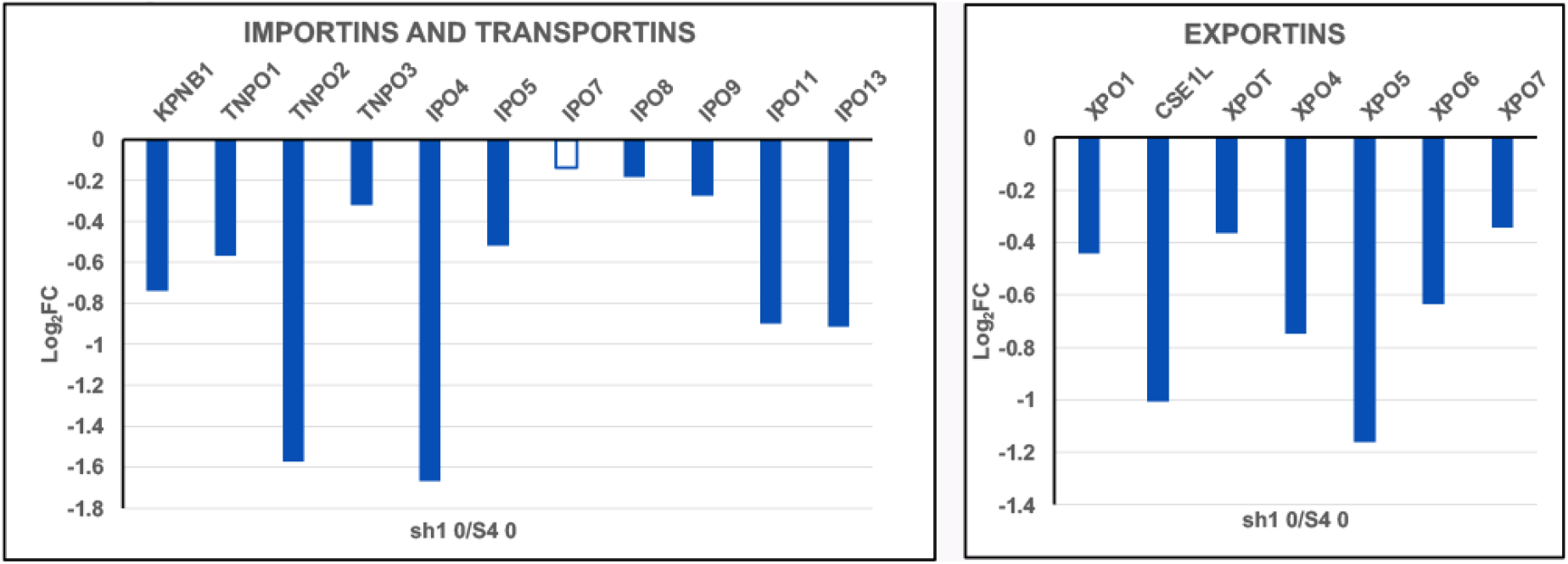
Importins, Transportins and Exportins relative transcript levels (Log_2_FC). Cell states: undifferentiated HL-60/sh1 0 cells compared to undifferentiated HL-60/S4 0 cells. The hollow bar indicates a lower statistical significance (PPDE <0.95); solid bars indicate a PPDE > 0.95.

**Figure 10.**
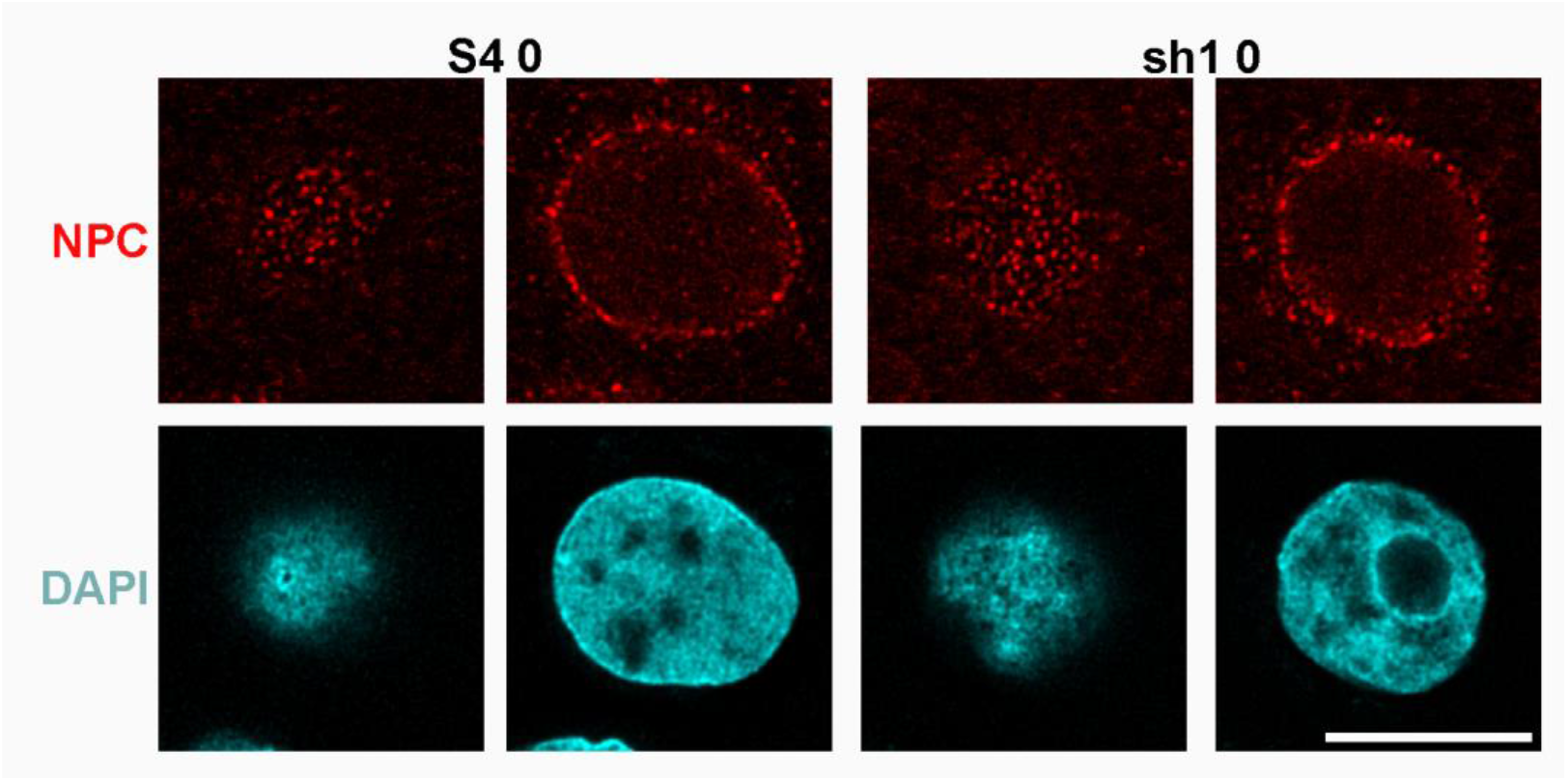
Immunostaining of an undifferentiated HL-60/S4 0 cell and an undifferentiated HL-60/sh1 0 cell with anti-nuclear pore complex (Mab 414) (Top Row), compared to the identical DAPI stained regions (Bottom Row). The first and third columns display tangential views of the cell nuclei; the second and fourth columns display mid-sections of the stained nuclei. All images are deconvolved. The magnification bar represents 10 μm.

It is striking that the Importin and Exportin transcript levels observed in sh1 0 cells, Figure 9, are downregulated, resembling the Importin and Exportin transcript situation with differentiated HL-60/S4, Figure 3.

GSEA was employed to examine the structure of nuclear pores in sh1 0 cells compared to control S4 0 cells (Figure 11) and Table 5. These analyses indicate enrichment of protein structure transcripts for the nuclear pores in sh1 0 cells. GSEA was also performed to determine whether the nuclear transport function is enriched in sh1 0 cells compared to control S4 0 cells (Figure 12) and Table 6. It is possible that the decreased levels of Importin and Exportin transcripts are not detrimental to the nuclear transport function. Perhaps, the upregulation of NUP protein transcripts can compensate for the decreased levels of Importin and Exportin transcripts.

**Table 5.** GSEA plot parameters from Figure 11. The positive NES values illustrate the relative enrichment of gene set transcripts in the LBR knockdown (sh1 0) phenotype region. The Nominal p values indicate that most of the NES scores and their structural implications are quite significant.

| GENE SET | Phenotypes | NES | Nominal p | Genes |
| --- | --- | --- | --- | --- |
| GOBP_NUCLEAR_PORE_ORGANIZATION | sh1 0/ S4 0 | 1.93 | 0.004 | 15 |
| GOBP_NUCLEAR_PORE_COMPLEX_ASSEMBLY | sh1 0/ S4 0 | 1.62 | 0.04 | 11 |
| GOCC_NUCLEAR_PORE_NUCLEAR_BASKET | sh1 0/ S4 0 | 1.67 | 0.01 | 13 |
| GOCC_NUCLEAR_PORE_OUTER_RING | sh1 0/ S4 0 | 1.58 | 0.03 | 10 |

**Table 6.** GSEA plot parameters from Figure 12. The positive NES values illustrate the relative enrichment of gene set transcripts in the LBR knockdown (sh1 0) phenotype region. The Nominal p values indicate that most of the NES scores (and their functional implications) are quite significant.

| GENE SET | Phenotypes | NES | Nominal p | Genes |
| --- | --- | --- | --- | --- |
| GOMF_NUCLEAR_EXPORT_SIGNAL_RECEPTOR_ACTIVITY | sh1 0/ S4 0 | 1.71 | 0.02 | 12 |
| GOMF_NUCLEAR_IMPORT_SIGNAL_RECEPTOR_ACTIVITY | sh1 0/ S4 0 | 1.52 | 0.04 | 20 |
| GOBP_NUCLEAR_EXPORT | sh1 0/ S4 0 | 1.66 | 0.0 | 169 |
| GOBP_NUCLEAR_TRANSPORT | sh1 0/ S4 0 | 1.60 | 0.0 | 338 |

**Figure 11.**
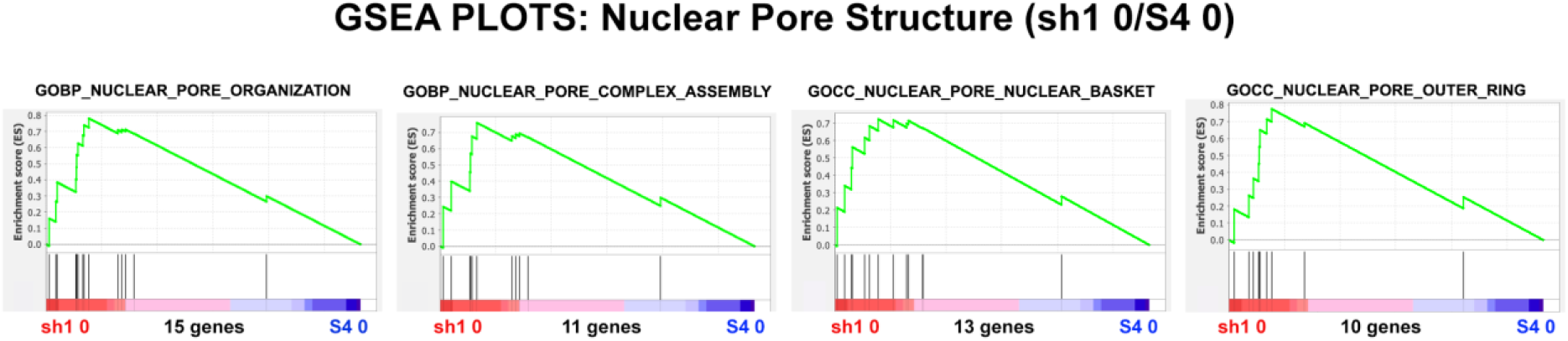
GSEA plots with four different nuclear pore structure gene sets. The number of genes in each gene set are shown at the bottom of each plot. The plots display undifferentiated (HL-60/sh1 0) cells compared to undifferentiated (HL-60/S4 0 cells). From the shape of these plots and the increased density of ranked genes at the sh1 0 phenotype location, it is clear that most of the genes of each gene set are over-represented in the LBR knockdown (sh1 0) phenotype region, implying that nuclear pore structure is **not** adversely affected by loss of LBR.

**Figure 12.**
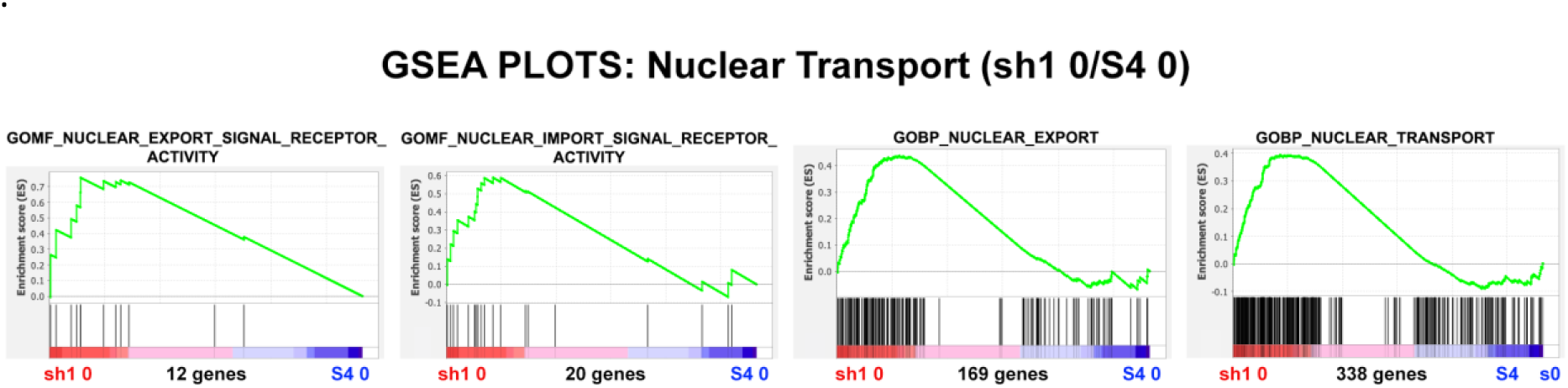
GSEA plots with four different nuclear transport gene sets. The number of genes in each gene set are shown at the bottom of each plot. The plots display undifferentiated HL-60/sh1 0 cells compared to undifferentiated HL-60/S4 0 cells. From the shape of these plots and the increased density of ranked genes at the sh1 0 phenotype location, it is clear that most of the genes of each gene set are over-represented in the LBR knockdown (sh1 0) phenotype region, implying that nuclear pore transport function is **not** adversely affected by loss of LBR.

**Figure 13.**
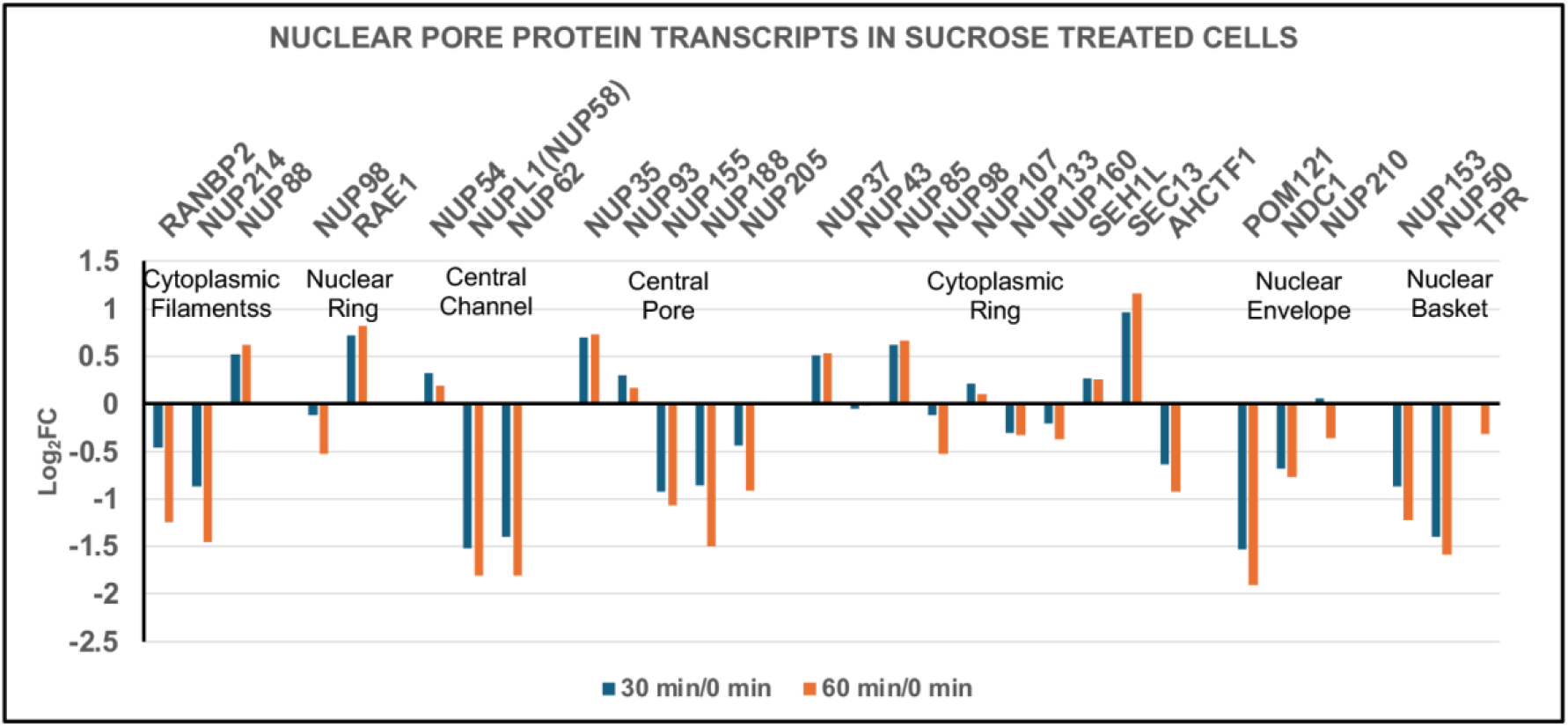
Nuclear pore protein transcript levels (Log_2_FC) during sucrose produced hyperosmotic cellular dehydration. Cell states: 30 min dehydration of undifferentiated HL-60/S4 cells compared to control undifferentiated HL-60/S4 cells (30 min/0 min); 60 min dehydration of undifferentiated HL-60/S4 cells compared to control undifferentiated HL-60/S4 cells (60 min/0 min). Seven NPC structural regions are indicated, with resident gene names clustered together.

In the present study, S4 0 cells served as a control for the experimental sh1 0 LBR knockdown cells. In our earlier study of LBR knockdown (Olins et al., 2010), we created a second control cell line (gfp 0), which employed a GFP sequence in a lentiviral vector, analogous to the sh1 0 construction, which employed a short hairpin “sh” RNA targeting LBR. Comparing the LBR protein levels of the three cell lines (S4 0, sh1 0 and gfp 0 by immunoblotting, clearly indicated that **only** the sh1 0 cells exhibited a significant reduction of LBR protein, see Figure 1 (Olins et al., 2010). GSEA comparisons of gfp 0/S4 0 transcripts have been obtained, analogous to Figures 11 and 12, and Tables 5 and 6. In all comparisons, the NES values for sh1 0/S4 0 were of greater magnitude than the NES values for gfp 0/S4 0. The sh1 0 cells grow robustly, similar to the S4 0 cells. Microscopic analyses show that the nuclei of sh1 0 cells are generally round-shaped, unlike the nuclei of S4 0 cells (Mark Welch et al., 2024; Olins et al., 2025b).

### Part III. The stress of cell dehydration

#### Hyperosmotic stressed undifferentiated HL-60/S4 cells exhibit disorganized changes in nuclear pore protein transcription

Exposure of HL-60/S4 to hyperosmotic medium (i.e., 300 mM sucrose in isosmotic tissue culture medium for 30 and 60 minutes) results in rapid dehydration of healthy growing cells, shrinkage of cell volumes to ∼2/3 of the initial volume, condensation (congelation) of interphase chromatin and mitotic chromosomes, “de-mixing” and “separation” of chromatin binding proteins, and cell death in a few days (Olins et al., 2020). Simultaneously, cellular metabolism starts moving into “high-gear”, with increased mitochondrial activity, increased ribosomal biosynthesis, increased proteasome activity, coupled with a reduction in heterochromatin and a decrease in mRNA synthesis (Olins et al., 2025a). We have described this new cellular metabolism as a “frantic attempt to rebuild cell structure in the face of inevitable death.” Ironically, from the perspective of nuclear pore protein transcription, activity is not readily interpretable. Figure 13 shows NUP protein transcript changes, comparing 30 and 60 minutes of dehydration to the initial time (0 minutes). Most NUP transcripts are downregulated, with a minority upregulated and very little change between 30 and 60 minutes. The presence of a normal Nuclear Basket and stable interactions with the nuclear envelope appear to be questionable. Figure 14 indicates that the majority of Importins and Exportins are significantly downregulated at 30 and 60 minutes.

**Figure 14.**
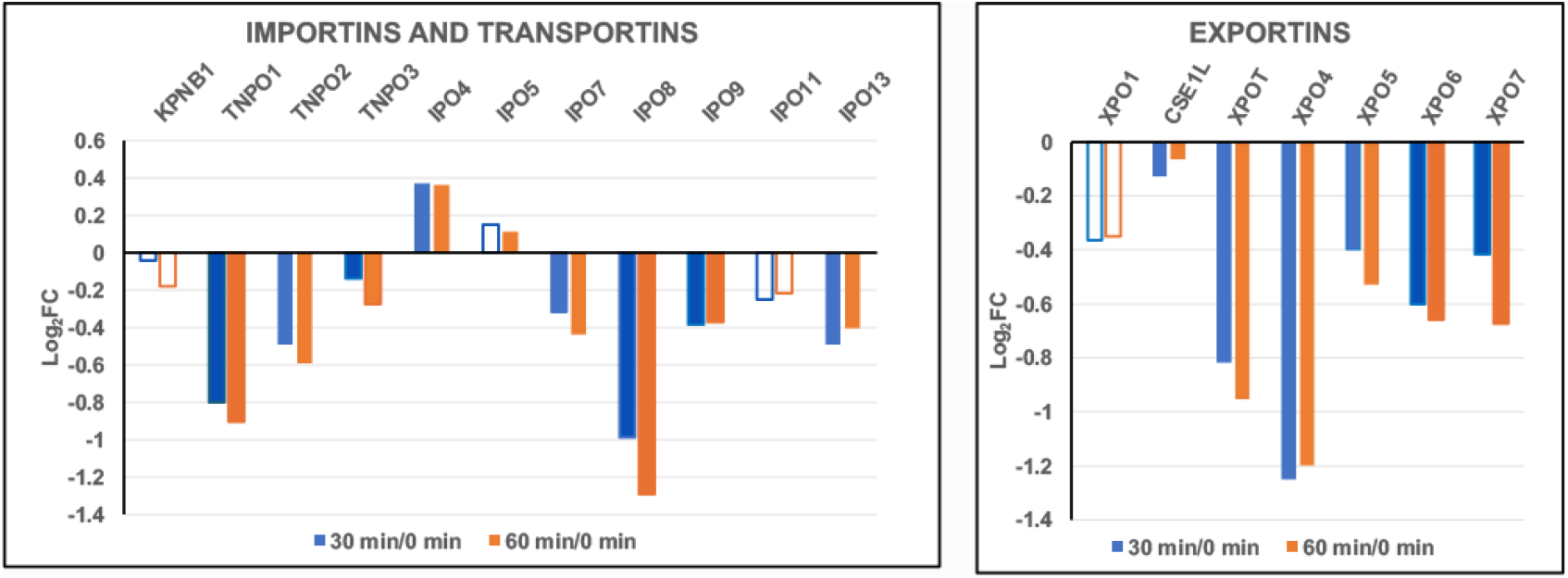
Importins, Transportins and Exportins transcript levels (Log2FC) during cell dehydration. Cell states: 30 min dehydration of HL-60/S4 cells, compared to control HL-60/S4 cells (30 min/0 min); 60 min dehydration of HL-60/S4 cells compared to control HL-60/S4 cells (60 min/0 min). Hollow bars indicate a lower statistical significance (PPDE <0.95); solid bars indicate a PPDE > 0.95.

Note that within a cluster, some genes are up and some are down. Note also, that for most genes, 30 and 60 minutes moved in the same direction.

GSEA Nuclear Pore Structure (Figure 15) indicates a general NPC disorganization and disassembly, but the Nominal p values are very poor, although improving between 30 and 60 minutes, see Table 7. GSEA Nuclear Transport (Figure 16) indicates a decrease in general export, import and transport, and in mRNA import. These various functions progressively deteriorate between 30 and 60 minutes, simultaneously as the Nominal p values improve in significance, see Table 8.

**Table 7.** GSEA plot parameters from Figure 15. The negative NES values indicate the relative paucity of gene set transcripts in the hyperosmotically-stressed cell (30 and 60 min) phenotype regions. The Nominal p values are very poor, although improving between 30 and 60 minutes.

| GENE SET | Phenotypes | NES | Nominal p | Genes |
| --- | --- | --- | --- | --- |
| GOBP_NUCLEAR_PORE_ORGANIZATION | 30 min/0 min | -0.90 | 0.59 | 15 |
|  | 60 min/0 min | -1.15 | 0.26 |  |
| GOBP_NUCLEAR_PORE_COMPLEX_ASSEMBLY | 30 min/0 min | -1.16 | 0.29 | 11 |
|  | 60 min/0 min | -1.32 | 0.13 |  |
| GOCC_NUCLEAR_PORE | 30 min/0 min | -0.85 | 0.81 | 95 |
|  | 60 min/0 min | -0.99 | 0.47 |  |

**Table 8.**
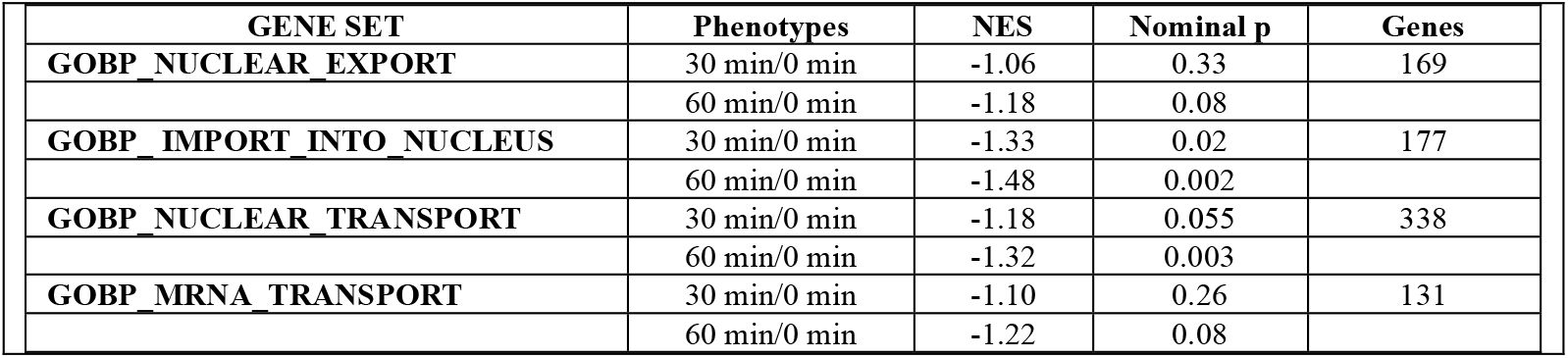
GSEA plot parameters from Figure 16. The negative NES values indicate the relative paucity of gene set transcripts in the hyperosmotically-stressed cell (30 and 60 min) phenotype regions. The Nominal p values are somewhat poor (although improving between 30 and 60 minutes).

| GENE SET | Phenotypes | NES | Nominal p | Genes |
| --- | --- | --- | --- | --- |
| GOBP_NUCLEAR_EXPORT | 30 min/0 min | -1.06 | 0.33 | 169 |
|  | 60 min/0 min | -1.18 | 0.08 |  |
| GOBP_IMPORT_INTO_NUCLEUS | 30 min/0 min | -1.33 | 0.02 | 177 |
|  | 60 min/0 min | -1.48 | 0.002 |  |
| GOBP_NUCLEAR_TRANSPORT | 30 min/0 min | -1.18 | 0.055 | 338 |
|  | 60 min/0 min | -1.32 | 0.003 |  |
| GOBP_MRNA_TRANSPORT | 30 min/0 min | -1.10 | 0.26 | 131 |
|  | 60 min/0 min | -1.22 | 0.08 |  |

**Figure 15.**
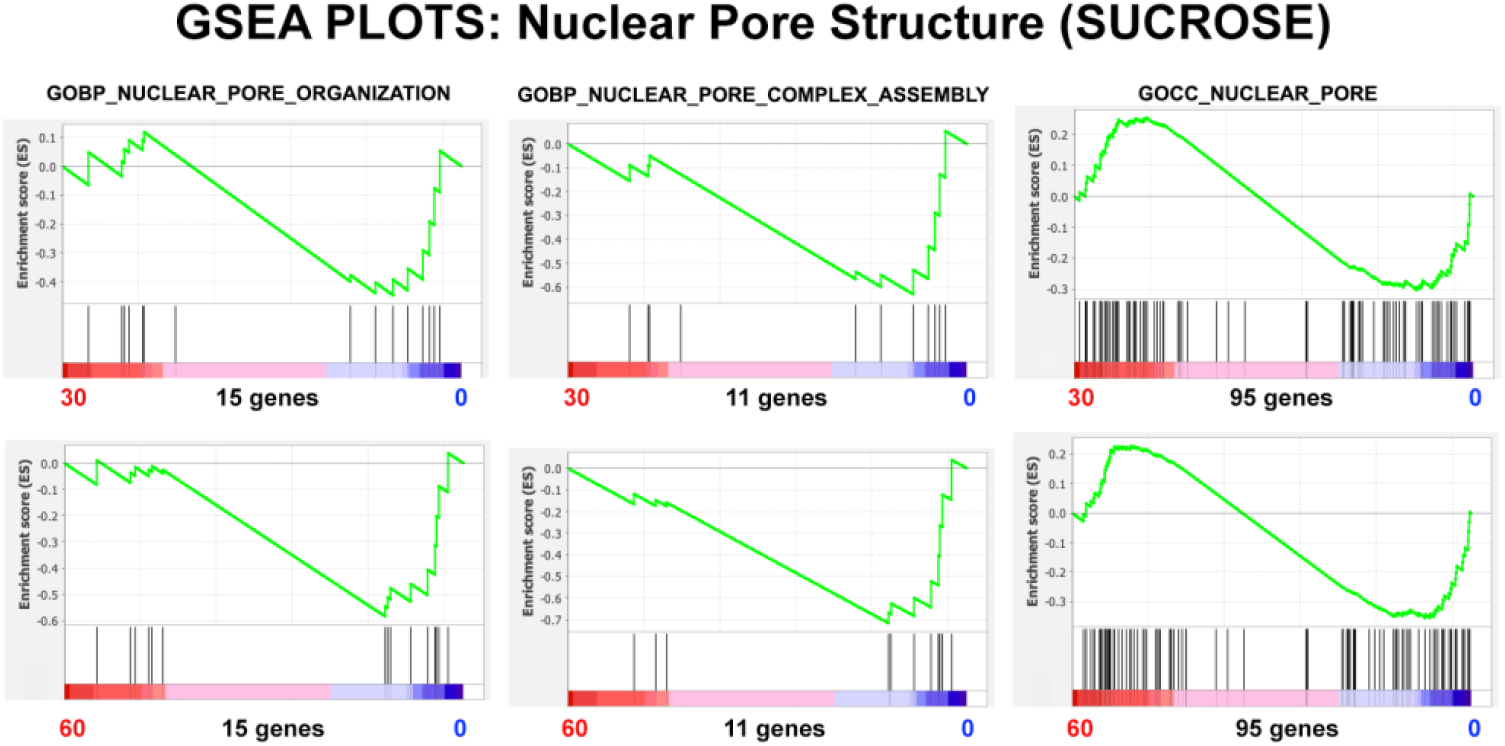
GSEA plots with three different nuclear pore structure gene sets. The number of genes in each gene set are shown at the bottom of each plot. The plots display HL-60/S4 0 cells exposed to hyperosmotic conditions (medium+300 mM sucrose) compared to control HL-60/S4 0 cells in isosmotic medium. Top row: 30 minutes of sucrose; Bottom Row, 60 minutes of sucrose. From the shape of these plots and the increased density of ranked genes at the control (0) phenotype location, it is clear that most of the genes in each gene set are underrepresented in the hyperosmotically-stressed cell (30 and 60 min) phenotype regions, implying that nuclear pore organization is not “normal” in the dehydrated state.

**Figure 16.**
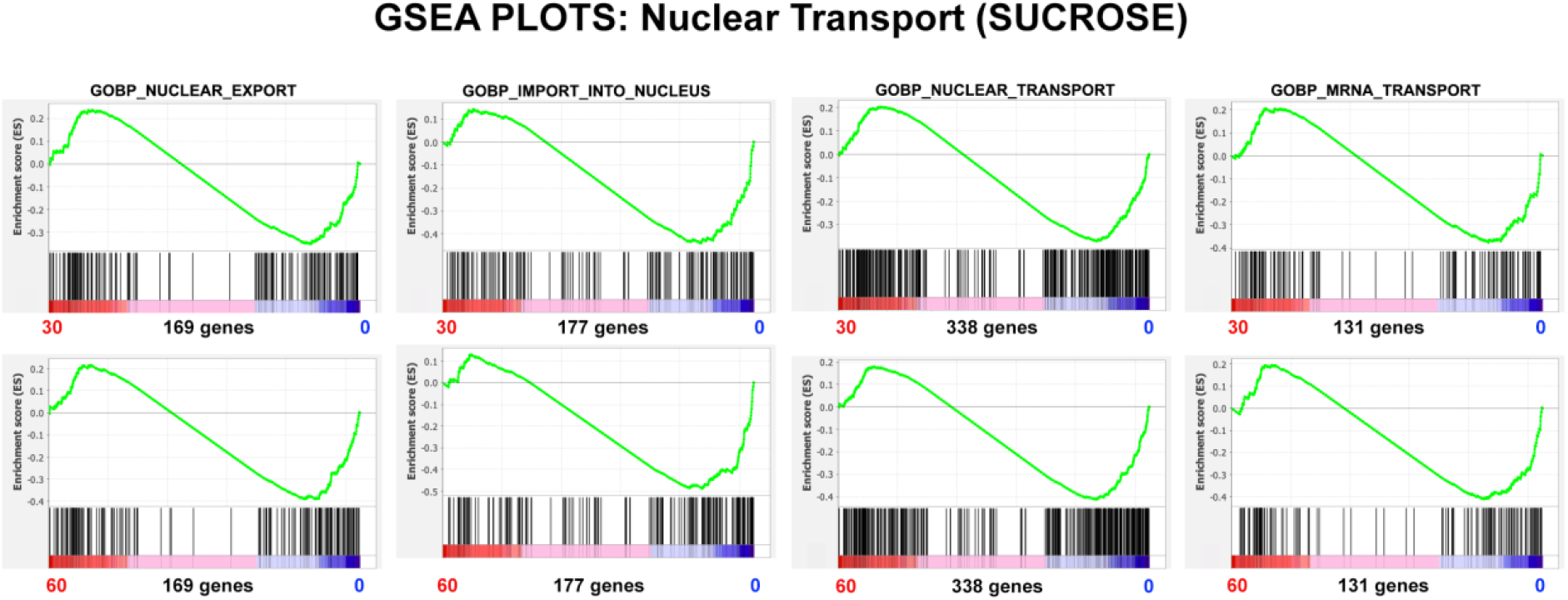
GSEA plots with four different nuclear transport gene sets. The number of genes in each gene set are shown at the bottom of each plot. The plots display HL-60/S4 0 cells exposed to hyperosmotic conditions (medium+300 mM sucrose) compared to control HL-60/S4 0 cells in isosmotic medium. Top row: 30 minutes of sucrose; Bottom Row, 60 minutes of sucrose. From the shape of these plots and the increased density of ranked genes at the control (0) phenotype location, it is clear that most of the genes in each gene set are underrepresented in the hyperosmotically stressed cells (30 and 60 min), implying that nuclear pore function is disabled in the dehydrated state.

The effect of hyperosmotic stress upon immunostaining is shown in Figure 17. Mab 414 appears to move into the perinuclear cytoplasm.

**Figure 17.**
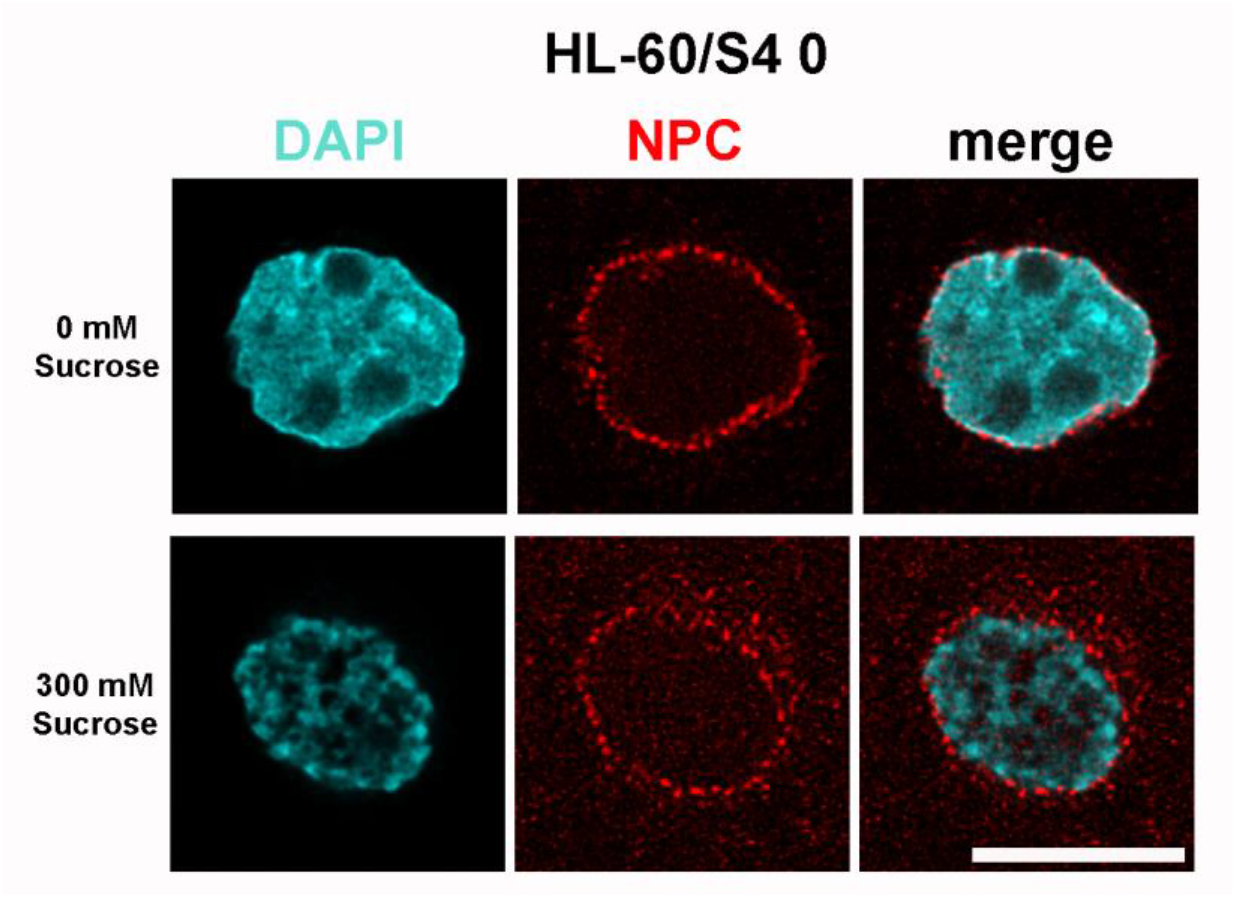
Immunostaining of HL-60/S4 0 using anti-nuclear pore complex (Mab 414), without sucrose dehydration (Top Row), compared to S4 0 cells exposed to 30 minutes of sucrose dehydration (Bottom Row). Note the congelation of interphase chromatin treated with sucrose, and the increased disorder of NPCs. All images are deconvolved. The magnification bar represents 10 μm.

## Discussion

Nuclear Pore Complexes (NPCs) are the principal conduits (channels) within the nuclear envelope, regulating the passage of small and large molecules to-and-from the nucleus and its surrounding cytoplasm. Excellent reviews have been published describing the structure of NPCs in mammalian cells (Buchwalter et al., 2019; Guglielmi et al., 2020; Han et al., 2022; Khan et al., 2020; Petrovic et al., 2025; Wu et al., 2025; Yang et al., 2023).

The purpose of the present article is to explore how the NPC responds, structurally and functionally, to various cellular stresses in a well-defined cell system. For this purpose, we have employed the myeloid leukemia cell line HL-60/S4 (Leung et al., 1992) which can be “stressed” in various ways: by induced cell differentiation (Mark Welch et al., 2017; Olins et al., 1998; Olins et al., 2001); by perturbation of the nuclear envelope, involving Lamin B Receptor (LBR) knockdown (Mark Welch et al., 2024; Olins et al., 2010; Olins et al., 2025b); and by cell and chromatin shrinkage resulting from hyperosmotic dehydration (Mark Welch et al., 2022; Olins et al., 2020; Olins et al., 2025a). Each of these three cellular “stresses” resulted in different combinations of nuclear pore structural and functional responses, illustrating the plasticity of Nuclear Pore Complexes.

### Part I. The stress of cell differentiation

Cell differentiation has been shown to be dependent upon transcript regulation of various NUPs. For example, an increase in transcript levels of NUP210 is required for myotube and neuronal differentiation in the mouse C2C12 myoblast cell line (D’Angelo et al., 2012). The opposite change of gene regulation has been observed in the differentiation of mouse cardiomyocytes (Han et al., 2022). The latter article argues: “The majority of Nup mRNAs decreased during cardiomyocyte maturation, with the exception of TPR and Nup45 (Figure 1G).” (Our Figure 2 displays some resemblance to their Figure 1G). The Han et al. article further concludes that the number and density of NPCs decrease during post-mitotic cardiomyocyte maturation. In HL-60/S4 cells, the apparent decreases in NPC structure and function parallel the short post-mitotic lives of RA-induced granulocytes and TPA-induced macrophage (Olins et al., 2024). These HL-60/S4 transcriptomes appear to focus upon their innate immunity functions (Mark Welch et al., 2017). It is interesting to note that in our Figures 2 and 3, the downregulation of NUP genes and of Importins and Exportins reveal considerable similarity, comparing the RA- and TPA-treated cells, possibly reflecting an overlap of their innate immunity functions.

### Part II. The loss of Lamin B Receptor

The increased majority of NUP protein transcripts in undifferentiated LBR knockdown cells (HL-60/sh1 0) compared to normal HL-60/S4 0 cells (Figure 8) is difficult to explain. Direct molecular interactions between LBR and NPC-related proteins are quite rare. One example has been described (Lu et al., 2010). The LBR N-terminal region was shown to interact with Importin beta (IPO5), which is downregulated in sh1 0 cells, compared to S4 0 cells (Figure 10). Likewise, the Importins and Exportins are generally downregulated in the undifferentiated sh1 0 cells. Possibly, their functions are not as important, given the significant increases in NUP protein transcripts. LBR is important for nuclear envelope growth, leading to nuclear lobulation and ELCS formation (Mark Welch et al., 2024; Olins et al., 2010; Olins et al., 2025b). In the HL-60/sh1 0 cells, the interphase nuclear shape appears more “round”, than in HL-60/S4 0 cells (Mark Welch et al., 2024; Olins et al., 2025b). The GSEA plot parameters (Tables 5 and 6), document enrichment of nuclear pore structure and nuclear transport in sh1 0 cells, compared to S4 0 cells. It is conceivable that the upregulated NUP transcripts in sh1 0 cells correlate with (and possibly produce) “rounder” and more functional (i.e., increased transport) sh1 0 nuclear envelopes. These, and other, structural and functional “advantages” of the sh1 0 cells versus the S4 0 cells have been recently described (Olins et al., 2025b). To quote our recent article: “We noted a significant increase in ribosomal protein and ribosomal RNA synthesis in sh1 0 cells, compared to the S4 0 cell line. Also, we noted a significant increase in heterochromatin and nucleosome formation in sh1 0 compared to S4 0 cells. This latter observation may be related to the dramatic morphological changes in DAPI staining of sh1 0 nuclei, which appeared to show chromatin condensates surrounding nucleoli.”

### Part III. The stress of cell dehydration

Hyperosmotic stress on HL-60/S4 0 cells produces dramatic changes in cell and nuclear shape, combined with congelation (condensation) of interphase and mitotic chromatin (Olins et al., 2020). CTCF and RAD21, which are believed to stabilize functional chromatin loops, are “de-mixed” to the exterior of congealed chromatin (Olins et al., 2020), suggesting a chaotic disorganization of gene regulation. The disorganization of gene expression resulting from hyperosmotic stress has been documented in additional studies (Mark Welch et al., 2022; Olins et al., 2025a). The present analyses demonstrate that hyperosmotically-stressed S4 0 cells exhibit poorly structured NPCs with functionally deficient nuclear transport (Tables 7 and 8), presumably contributing to their inevitable death.

#### Summarizing the consequences of the above selected stresses upon the NPCs and subsequent cell fates

In the present article, it is surprising that the undifferentiated LBR knockdown cells (HL-60/sh1 0) suffer only minor consequences to the growth and health of these cells. The NPCs of HL-60/sh1 0 cells exhibit “enriched” structure and transport function (Figures 11 and 12; Tables 5 and 6), when cultivated in their normal growth medium. However, RA treatment of HL-60/S4, HL-60/sh1 and HL-60/gfp cells result in eventual death by apoptosis (Olins et al., 2025b). The stresses of cell differentiation (Figures 4 and 5; Tables 3 and 4) may contribute to the eventual deaths of these three cell lines. Cellular dehydration from hyperosmotic sucrose treatment also results in rapid cell death, likely an indication of nuclear pore destruction and functional failure (Figures 15 and 16; Tables 7 and 8). The hyperosmotically stressed cells appear disorganized in their homeostatic response, a likely mechanism for inevitable death.

## Acknowledgements

ALO and DEO express our gratitude to the MaineHealth Institute for Research for allowing us to join the research group of Dr. Igor Prudovsky. We thank David Mark Welch for his analysis of the transcriptome data and for formatting the data for GSEA. Confocal microscopy studies were performed using a Leica SP-8 microscope at the Histopathology and Microscopy core (supported by NIH COBRE grant P30GM106391) of MaineHealth Institute for Research.

## Supplemental Files

**Table S1** GSEA Cell Differentiation Data Set

**Table S2** GSEA Loss of LBR Data Set

**Table S3** GSEA Cell Dehydration Data Set

